# Angiotensin-converting enzyme 2, a SARS-CoV-2 receptor, is upregulated by interleukin-6 via STAT3 signaling in rheumatoid synovium

**DOI:** 10.1101/2020.05.26.115261

**Authors:** Sho Mokuda, Tadahiro Tokunaga, Junya Masumoto, Eiji Sugiyama

## Abstract

Detected in December 2019, the coronavirus disease 2019 (COVID-19) has since spread all over the world, resulting in a global pandemic. The disease is caused by severe acute respiratory syndrome-coronavirus-2 (SARS-CoV-2), and its symptoms usually include cough, fever, and gastrointestinal problems. Although the prevalence of rheumatoid arthritis (RA) is about 1 % of the global population and RA patients naturally have a chance of acquiring COVID-19 in this pandemic, no studies have considered the expression of angiotensin-converting enzyme 2 (ACE2) (a receptor for SARS-CoV-2) in synovial tissues. Our presenting data revealed that ACE2 expression was elevated in active rheumatoid synovium, and siRNA against STAT3 was able to downregulate ACE2 expression, which was in turn induced by IL-6 signaling.

## Letters

Severe acute respiratory syndrome-coronavirus-2 (SARS-CoV-2) has spread explosively worldwide and resulted in a pandemic of a new respiratory disease called coronavirus disease 2019 (COVID-19) in March 2020. Symptoms of COVID-19 include fever, malaise, cough, and in severe cases, pneumonia and acute respiratory distress syndrome (1, 2). Coronavirus possesses envelope-anchored spike proteins that bind host cell surface receptors and allow viral entry into target cells. In the case of SARS-CoV-2, its spike protein mediates binding of the angiotensin-converting enzyme 2 (ACE2) receptor (3). While ACE2 expression levels are relatively low in the airway mucosa and lungs, this protein is predominant in the stomach, intestines, gallbladder, kidney, and heart (4). Therefore, in COVID-19 patients, SARS-CoV-2 can be detected not only in nasal and oral swabs but also in rectal samples; viral RNA has also been detected in patient blood (5).

Rheumatoid arthritis (RA) is an autoimmune disease that is characterized by systemic synovitis and affects about 1% of the world population. The expression pattern of ACE2 in synovial tissues has not yet been reported, although RA patients naturally have a chance of getting COVID-19 in this pandemic (6–8). In this study, we investigated ACE2 expression in the synovium and its regulatory expression mechanism. Immunohistochemistry analysis revealed that the active rheumatoid synovium, where remarkable thickening of the synovial lining and mesenchymoid transformation of the synovial stroma were observed, displayed a higher expression of ACE2 as compared to inactive samples (Figure 1A). Reverse transcriptase-quantitative polymerase chain reaction (RT-qPCR) for ACE2 mRNA extracted from the synovium supported these data (Figure 1B). ACE2 expression was found to be increased in the synovial lining and sublining regions, suggesting that its expression was elevated in fibroblast-like synoviocytes (FLS). Primary cultures of these RA-derived FLS revealed that interleukin-6 (IL-6) stimulation increased ACE2 expression (Figure 1C). IL-6 is known to regulate downstream target genes via signal transducer and activator of transcription 3 (STAT3) signaling activation (9). IL-6 stimulation led to tyrosine phosphorylation of STAT3 (data not shown). Indeed, the use of small interfering RNA (siRNA) against STAT3 reduced the IL-6-dependent ACE2 expression in RA-FLS (Figure 1D).

**Figure 1.**
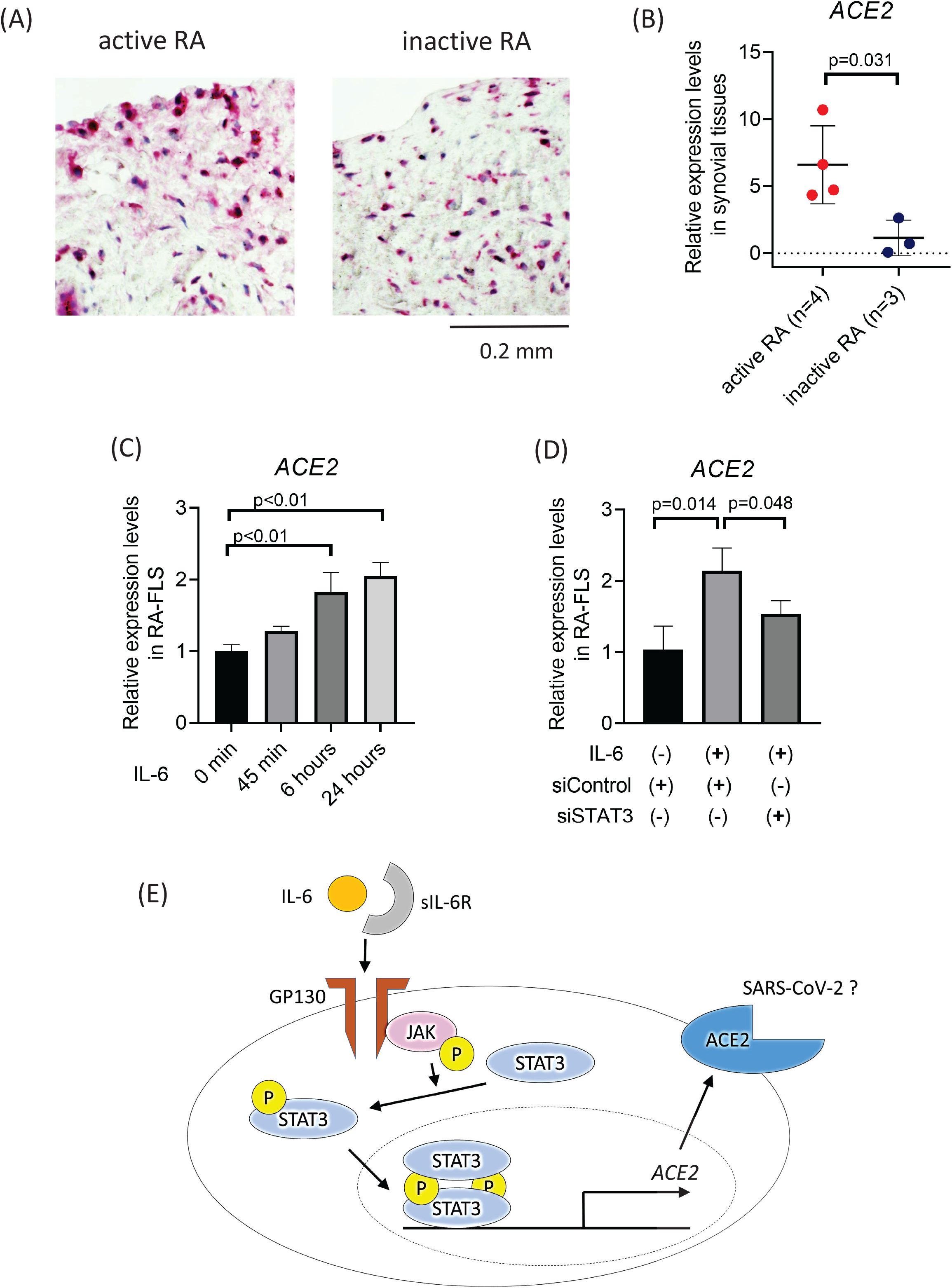
ACE2 upregulation in inflamed synovial tissues via IL-6 signaling. (A) Immunohistochemistry for ACE2 in rheumatoid synovial tissues. ACE2 was stained with Vector Red (red). Images are representative data of 3 specimens. Bar = 0.2 mm. (B) RT-qPCR for ACE2 expression levels in rheumatoid synovial tissues (n = 3–4). Data are presented as means ± SD. Active rheumatoid synovitis was defined by the characteristics of multi-layered lining (hyperplasia), mesenchymoid transformation, palisading appearance of synovial lining and proliferation of blood vessels, all detected with hematoxylin and eosin (H & E) staining. (C) RT-qPCR for ACE2 expression levels in RA-FLS stimulated with rIL-6 (100 ng/ml) and rIL-6Rα (100 ng/ml) (n = 3). Data are presented as means ± SEM. (D) RT-qPCR (mean ± SEM, n = 3) for ACE2 expression levels in RA-FLS stimulated with rIL-6 (100 ng/ml), rIL-6Rα (100 ng/ml), and siRNA against STAT3. (E) Schematic representation of IL-6-STAT3 signaling-mediated ACE2 expression. Expression levels detected with RT-qPCR were normalized to GAPDH expression. Data were analyzed using a t-test.

We propose that ACE2 is upregulated in the synovium of active RA patients and is likely maintained by STAT3-mediated activation through the IL-6 pathway (Figure 1E). Therefore, RA activity may be able to alter SARS-CoV-2 entry into synovial cells. Further analyses of IL-6-induced ACE2 transcription in cells other than FLS may help elucidate the mechanisms of viral entry into target cells, as *in vivo* IL-6 levels are high in COVID-19 patients (2). These data also implies that unplanned preventive withdrawal of disease-modified antirheumatic drugs (DMARDs) could lead to increase the risk of COVID-19.

## Methods

### Study design

This study was approved by the clinical ethics committees of Hiroshima University Hospital, Dohgo Spa Hospital and Ehime University Proteo-Science Center and Graduate School of Medicine and was conducted at these institutions (approval number: E-668; approval date: 01/02/2017). All experiments were performed in accordance with the approved guidelines. Synovial tissues were collected from 16 rheumatoid arthritis (RA) patients that fulfilled the classification criteria of the 1987 American College of Rheumatology (Arthritis and Rheum. 1988; 31: 315–324) and underwent total joint replacement, after obtaining informed and signed consent forms. Patients were not involved in the design, conduct, reporting, or dissemination plans of our research. Active rheumatoid synovitis was defined by the characteristics of multi-layered lining (hyperplasia), mesenchymoid transformation, palisading appearance of synovial lining and proliferation of blood vessels, all detected with hematoxylin and eosin (H & E) staining.

### RNA extraction and reverse transcription quantitative PCR (RT-qPCR)

Total RNA was extracted and purified from synovial tissue (a total of 7 samples from 7 different RA patients) and cultured cells using Trizol reagent (Life Technologies, Carlsbad, CA, USA), followed by cDNA synthesis using a PrimeScript RT Reagent Kit with gDNA Eraser (Takara-Bio, Kusatsu, JAPAN). RT-qPCR using Brilliant II SYBR® Green QPCR Master Mix (Agilent, Santa Clara, CA, USA) was outperformed on CFX Connect Real-Time PCR Detection System (Bio-Rad Laboratories, Hercules, CA, USA). The upstream and downstream primer sequences for angiotensin-converting enzyme 2 (ACE2) were 5′-GATTCTTTTTGGGGAGGAGGA-3′ and 5′-CTCCGGGACATCCTGATG-3′, respectively. The upstream and downstream primer sequences for human glyceraldehyde 3-phosphate dehydrogenase (GAPDH) were 5′-AAGGTCATCCCAGAGCTGAA-3′ and 5′-CTGCTTCACCACCTTCTTGA-3′, respectively.

### Immunohistochemistry

Sections obtained from formalin-fixed paraffin-embedded tissues (a total of 6 samples from 6 different RA patients) were used for immunohistochemistry. Following deparaffinization and antigen retrieval in Tris-EDTA buffer (pH 9.0), the specimens were first incubated with primary antibodies (anti-ACE2, rabbit polyclonal, Bioss Antibodies Inc., Woburn, MA, USA), and then with the AP-conjugated secondary antibodies. The samples were then visualized using ImmPACT Vector Red Alkaline Phosphatase Substrate (Vector Laboratories, Burlingame, CA, USA), and the slides were counterstained with hematoxylin solution.

### Purification and culture of fibroblast like synoviocytes (FLS)

To obtain FLS, synovial tissues from three different RA patients were minced and incubated with 1mg/ml collagenase/dispase (Sigma-Aldrich, Tokyo, JAPAN) in PBS (pH7.2) for 1 hour at 37 °C. The tissue samples were then filtered, washed, and cultured. During the culture, the supernatant was replaced frequently to remove nonadherent cells. The FLS were cultured in DMEM (FUJIFILM Wako Pure Chemical Corporation, Osaka, JAPAN) supplemented with 10% fetal bovine serum (Sigma-Aldrich) and penicillin/streptomycin (FUJIFILM Wako Pure Chemical Corporation). Cultured cells at passages 3 through 6 were used for the experiments and were starved in DMEM with 0.5% FBS for > 6 before the addition of recombinant cytokines, human Interlukin-6 (IL-6) (100 ng/ml) and human IL-6Rα (100 ng/ml) (BioLegend, San Diego, CA, USA).

### Transfection experiments using siRNA

A total of 3 × 10^4^ FLS per well in 12-well plates were transfected with siRNA at 20 pmol using ScreenFect siRNA (ScreenFect GmbH, Eggenstein-Leopoldshafen, Germany), according to the manufacturer’s protocol. The siRNA targeting STAT3 (FlexiTube siRNA, SI02662898) and negative control (Allstar Negative Control siRNA) were purchased from QIAGEN (Shanghai, China).

### Statistical analysis

All graphs show the results of one representative experiment from the several individual experiments performed. All statistical analyses were performed using Student’s t-test. Data were processed and analyzed using the GraphPad Prism 8 software (Graph Pad Software Inc., La Jolla, CA, USA).

## Acknowledgements

We thank the staffs of the Department of Clinical Immunology and Rheumatology at Hiroshima University Hospital, Japan, and the staffs of Dohgo Spa Hospital, Matsuyama, Ehime, Japan, for the preparation of patients’ specimens.

## Notes

**Conflict of Interest**: None.

### Competing Interest Statement

The authors have declared no competing interest.

